# Kidney-specific methylation patterns correlate with kidney function and are lost upon kidney disease progression

**DOI:** 10.1101/2022.09.19.508466

**Authors:** Naor Sagy, Noa Meyrom, Pazit Beckerman, Oren Pleniceanu, Daniel Z Bar

## Abstract

Chronological and biological age correlate with DNA methylation levels at specific sites in the genome. Linear combinations of multiple methylation sites, termed epigenetic clocks, can inform us of the chronological age and predict multiple health-related outcomes. However, why some sites correlate with lifespan, healthspan, or specific medical conditions remains poorly understood. Kidney fibrosis is the common pathway for Chronic Kidney Disease, which affects 10% of Europe and USA population. Here we identify epigenetic clocks and methylation sites that correlate with kidney function. Moreover, we identify methylation sites that have a unique methylation signature in the kidney. Methylation levels in the majority of these sites correlates with kidney state and function. When kidney function deteriorates, all of these sites regress towards the common methylation pattern observed in other tissues. Interestingly, while the majority of sites are less methylated in the kidney and become more methylated with loss of function, a fraction of the sites are highly methylated in the kidney and become less methylated when kidney function declines. These methylation sites are enriched for specific transcription-factor binding sites. In a large subset of sites, changes in methylation pattern are accompanied by changes in gene expression in kidneys of chronic kidney disease patients. These results support the information theory of aging, and the hypothesis that the unique tissue identity, as captured by methylation patterns, is lost as tissue function declines. However, this information loss is not random, but guided towards a baseline that is dependent on the genomic loci.

## Introduction

### Chronic kidney disease (CKD)

The human kidney plays several essential roles, including the excretion of toxic waste products and maintenance of blood pressure and pH. The average kidney contains ~1 million nephrons, which are its main functional units (Fairweather et al. 2020). CKD is defined as loss of renal function, marked by decreased glomerular filtration rate (GFR)<60 mL/min/1.73m^2^, and increase in kidney damage markers (e.g., albuminuria), or both, for ≥3 months (Anon 2013). Renal function can be assessed using serum creatinine (sCr) levels, used to compute the estimated Glomerular Filtration Rate (eGFR) via one of several formulae (Musso et al. 2016). Unfortunately, sCr-based assessments are highly inaccurate due to several reasons: (1) sCr is highly insensitive for diagnosing early CKD, as abnormalities in its values occur only when ~40% of the renal parenchyma is damaged (Steubl et al. 2016). Indeed, many individuals with normal sCr actually have kidney damage upon histological examination including interstitial fibrosis (IF), tubular atrophy (TA) and glomerulosclerosis (GS) - all indicative of impaired renal function (Mancilla et al. 2008; Okamoto et al. 2010; El Agha et al. 2017). CKD is a relentless disorder, steadily deteriorating from its early stages (1-2) in which the GFR is normal or high to stage 5, or end-stage kidney disease (ESKD), defined as a GFR<15 ml/min/1.73m2, when patients require kidney replacement therapy (KRT): either dialysis or transplantation (Anon 2013). With a prevalence of up to 17.3% (Brück et al. 2016) requiring expensive treatments, CKD is a global epidemic, accounting for over million deaths a year (GBD 2015 Mortality and Causes of Death Collaborators 2016). Moreover, its true prevalence is hard to determine, since its early stages often go undetected. CKD rates are expected to keep rising due to the aging of the population and growing prevalence of its 2 main risk factors: diabetes mellitus (DM) and hypertension (HTN) (GBD 2015 Mortality and Causes of Death Collaborators 2016). The irreversible nature and lack of specific treatments for CKD underscore the importance of its early diagnosis, while renal function is preserved and active interventions (e.g., glycemic control in DM) may slow its progression, which will improve quality of life, reduce mortality, and reduce the costs for healthcare systems (Levin & Stevens 2011).

### DNA methylation correlates with lifespan, healthspan and CKD

DNA methylation is an epigenetic modification, and the most studied one in relation to diseases, healthspan and lifespan. Seminal works by Horvath and Hannum described ‘epigenetic clocks’, a linear combination of the methylation levels of several tens to hundreds of genomic sites, which accurately estimate the biological age of most human tissues, including the kidney (Lu et al. 2019; Horvath 2013; Hannum et al. 2013). These clocks have been expanded to predict “biological age”, capture multiple aspects of healthspan and predict all-cause mortality (Lu et al. 2019). Proving the relevance of epigenetic signatures to renal function, the Susztak group profiled genome-wide methylation of tubular cells in CKD and normal kidneys and found that a 65-probe signature correlates with kidney structural damage, as seen in biopsy, and a 471-probe signature predicts renal functional decline together with clinical parameters (Gluck et al. 2019). Likewise, it has been recently shown that the progression of CKD is tightly linked to methylation-induced changes in tubular cell function, including senescence (Al-Dabet et al. 2022). Remarkably, renal methylation signatures also accurately reflect the presence of kidney cancer, with the methylome of RCC cells exhibiting a significantly older biological age, a phenomena seen in a wide range of cancers. Moreover, it has been shown that specific DNA methylation signatures predict various clinical outcomes, and correlate with renal function (Lu et al. 2019; Dritsoula et al. 2021; Rysz et al. 2022; Schlosser et al. 2021).

### Changes in DNA methylation alter transcription factor binding

DNA methylation often correlates with transcription levels. Epigenome-wide association studies (EWAS) demonstrated that methylation at specific CpGs sites are associated with kidney disease (Wing et al. 2014; Chu et al. 2017; Ko et al. 2013; Sapienza et al. 2011; Smyth et al. 2014; Rakyan et al. 2011). Methylation and demethylation can have direct effects on expression levels *in vivo* (Kangaspeska et al. 2008; Métivier et al. 2008) thus suggesting that some of the correlations might have a causal component to them. Methylation signatures can alter some TFs binding (Bonder et al. 2017; Héberlé & Bardet 2019), which can mechanistically explain some of this effect.

Here, we identify epigenetic clocks and CpG sites that correlate with the functional state of the kidney, as captured by IF and eGFR. A subset of these CpG sites show a kidney specific methylation pattern. Moveover, the majority of sites showing a kidney specific methylation pattern correlate with IF. Multiple genes associated with these sites are changed in CKD. Interestingly, all these sites lose their unique methylation pattern in CKD, while trending towards the common form.

## Results

### Specific epigenetics clocks correlate with eGFR and IF

CKD progression is accompanied by a decrease in eGFR and an increase in IF (Mancilla et al. 2008; Okamoto et al. 2010; El Agha et al. 2017). Several studies have analyzed the methylation of DNA in kidney tissue and identified CpGs whose methylation levels correlate with these markers of kidney disease progression (Schlosser et al. 2021; Ko et al. 2013; Bontha et al. 2017). To test whether eGFR decrease and IF increase are captured by existing epigenetic DNA methylation clocks, we analyzed 85 publicly available kidney methylation arrays (Ko et al. 2013; Bontha et al. 2017). Multiple clocks showed a significant correlation with eGFR and IF (**Supp. Fig. 1**; **Supp. Table 1**). We noticed substantial differences in specific clocks performed between males and females, thus we performed these analyses independently (**Fig. 1,2**; **Supp. Tables 2-3**). Of all clocks tested, DNAmB2M performed best in males, giving a correlation of −0.55 (p=4**×** 10^−4^) to eGFR and 0.75 (p=7**×** 10^−8^) to IF. By contrast, in females, DNAmPhenoAge gave a correlation of −0.39 (p=0.017) to eGFR and DNAmPAI1 of −0.39 (p=0.015) to IF. Age and BMI, two potential confounders, were poorly correlated with these clocks (**Supp. Tables 2-3**). We concluded that some aspects of the physiological state of the kidney are reflected in the methylome and captured by epigenetic clocks not specifically developed for this purpose.

**Figure 1:**
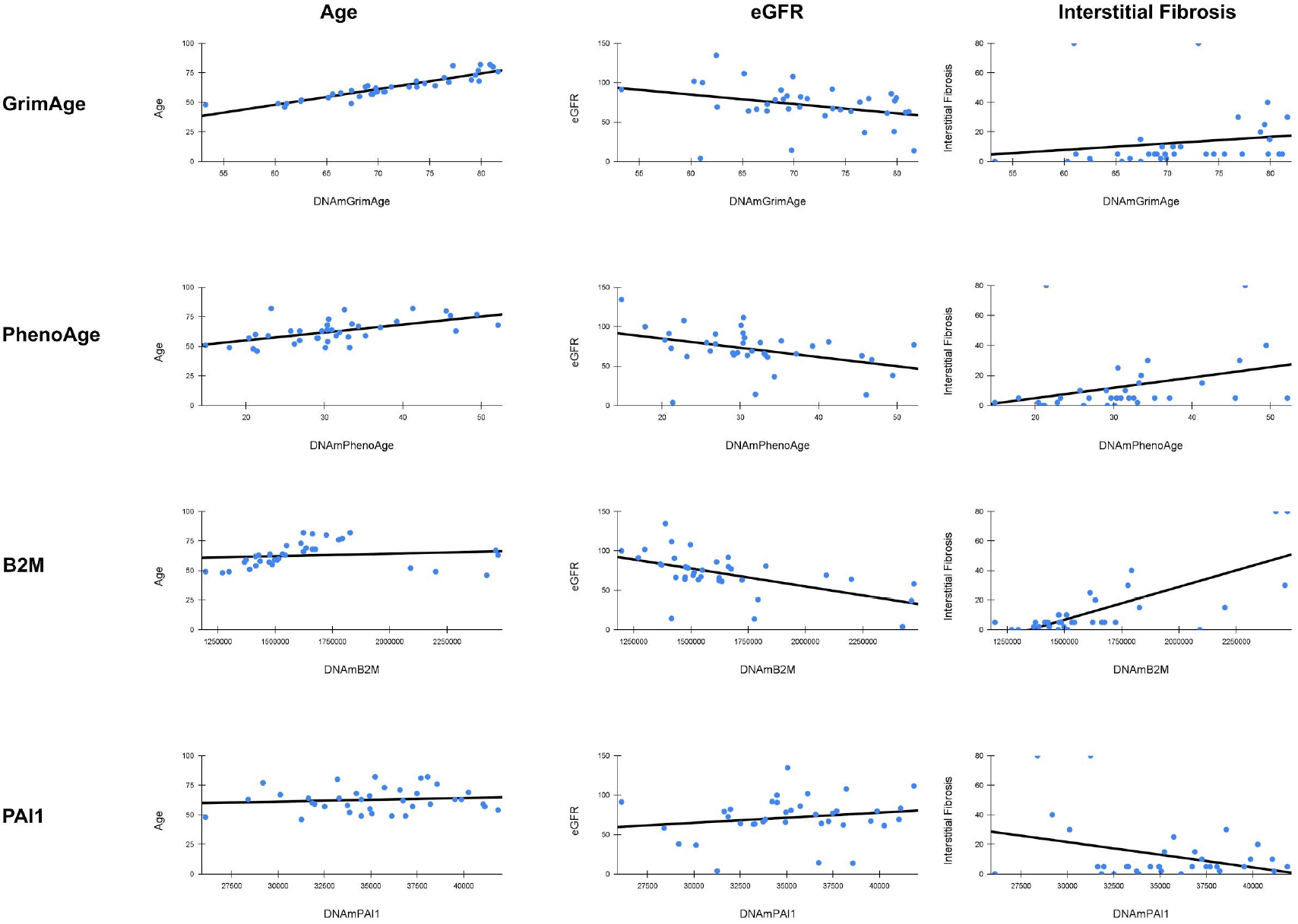
Methylation clocks correlate with kidney state in males. Correlation of age, eGFR and IF with four epigenetic clocks: GrimAge, PhenoAge, B2M and PAI1.

**Figure 2:**
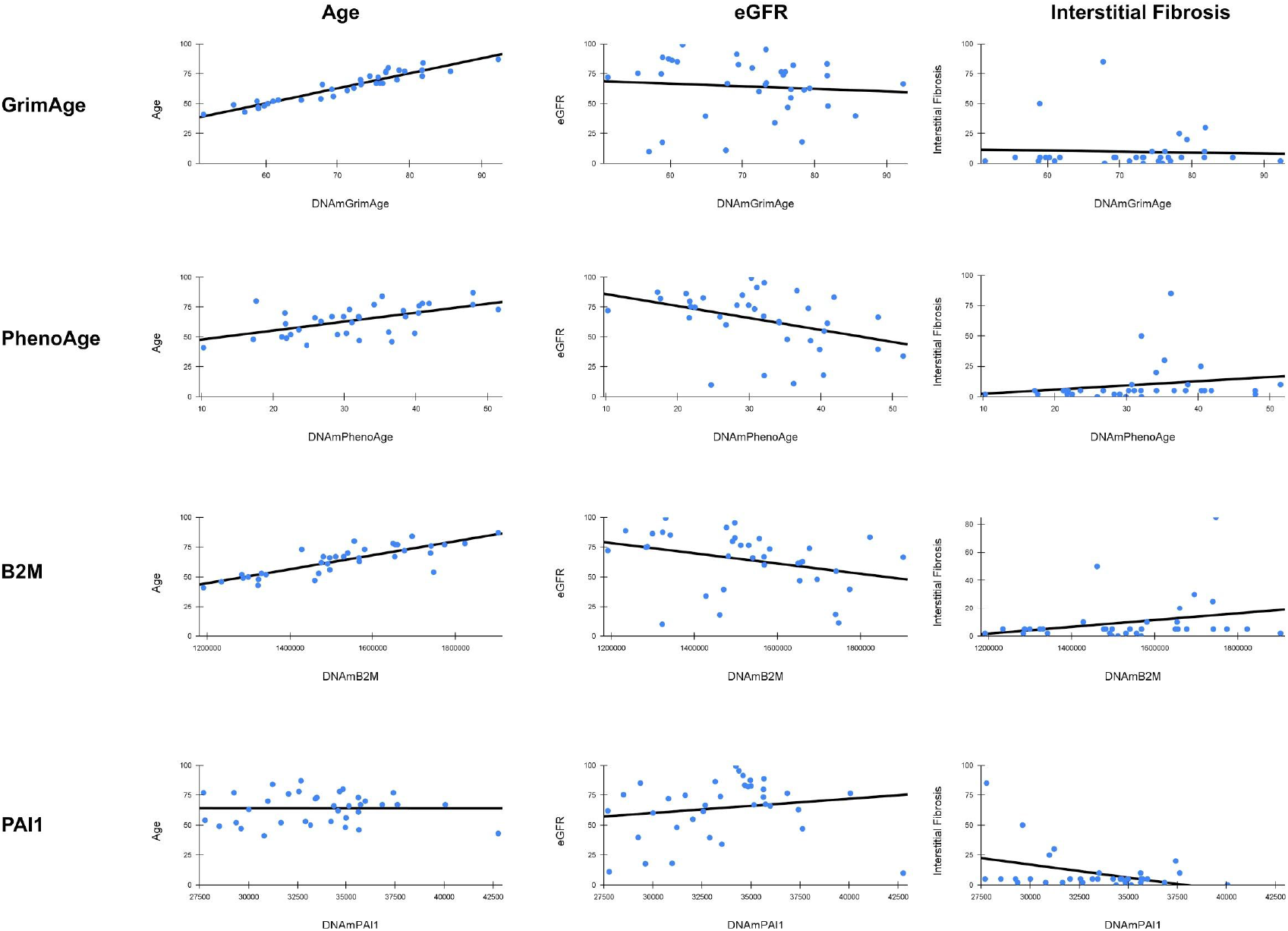
Methylation clocks correlate with kidney state in females. Correlation of age, eGFR and IF with four epigenetic clocks: GrimAge, PhenoAge, B2M and PAI1.

### Multiple CpG sites show strong correlation to eGFR and IF

Alternative methylation sites may provide an even better estimate of tissue state. Despite the small number of arrays, 1975 sites showed a statistically significant correction (after Bonferroni correction) for eGFR, with correlation values up to r=0.74 (**Fig. 3A,B**, **Supp. Table 4**). We tested if these results can be explained by simple global loss or gain of methylation. Of the 1975 sites, 42% positively and 58% negatively correlated with eGFR, excluding global unidirectional changes as drivers of these methylation patterns. For IF, 20,986 sites passed Bonferroni correction, with r as high as 0.89 (**Fig. 3C,D**, **Supp. Table 5**). Of these, 67% showed a positive correlation and 33% a negative correlation. To test for the possibility that age, sex, race or BMI act as confounders, we measured their correlation eGFR and IF in this dataset. Additionally, we examined the correlation of top CpG sites with age and BMI in this and the independent NGDC-CNCB (Xiong et al. 2021) dataset. In all cases, correlation to age, sex and BMI was weak or negligible, excluding them as confounders (**Supp. Table 6**). We took a closer look at cg10832035, the site with the highest correlation to eGFR (r=−0.74; p<10^−13^). As expected, it also showed a strong correlation to IF (r=0.77, p<10^−14^). It was located on a CpG island inside the non-coding gene RP5-1086L22.1. The adjacent methylation sites within the island, but not outside the island, exhibited a similar methylation pattern and correlation to IF (r=0.7; p<10^−10^) and eGFR (r=−0.64; p<10^−8^), excluding possible probe issues. Interestingly, this site was almost completely methylated in all tissues except the kidney (**Fig. 3E**). In the kidney, methylation levels averaged around 0.35, and tended to increase with IF and decline in eGFR.

**Figure 3:**
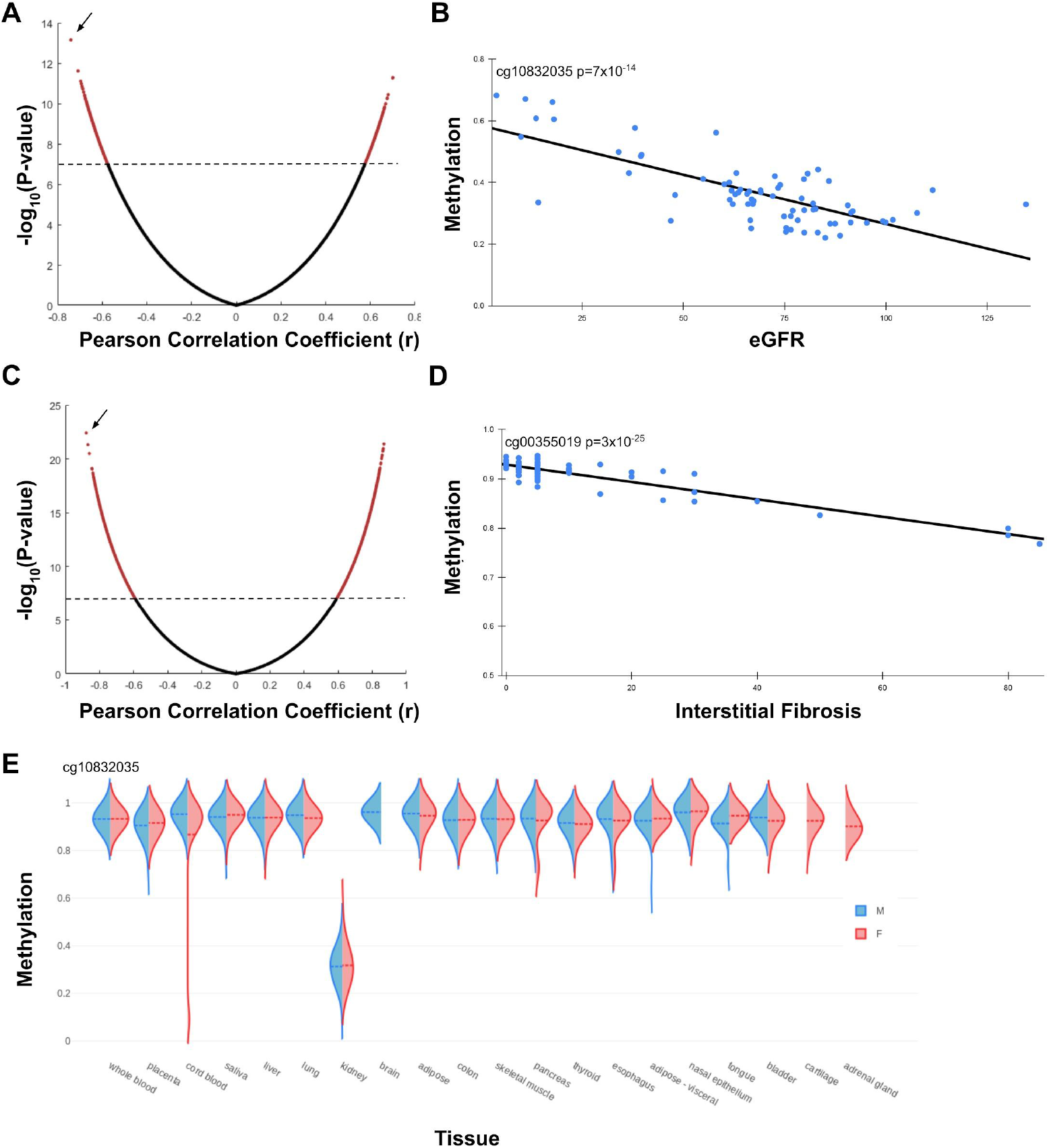
Individual CpG sites correlate with kidney state. **A**. Volcano plot of all tested CpG sites and their correlation to eGFR. Dots above the dashed line (red) pass Bonferroni correction. **B**. Individual methylation and eGFR levels for cg10832035, the site with highest correlation to eGFR (arrow in A). **C**. Volcano plot of all tested CpG sites and their correlation to IF. **D**. Individual methylation and eGFR levels for cg00355019, the site with highest correlation to IF (arrow in C). **E**. cg10832035 methylation patterns in a subset of tissues from the NGDC-CNCB dataset.

### CKD-correlated CpG sites are enriched for TF binding sites

To test for potential biological significance, sequences surrounding these CKD-correlated sites were used for motif enrichment analysis. Indeed, these sequences were enriched for transcription factor (TF) binding sites. Specifically, sites that positively correlated with IF showed enrichment for the HNF1 family, while negatively correlated sites were enriched for Jun/FOS (**Fig. 4**).

**Figure 4:**
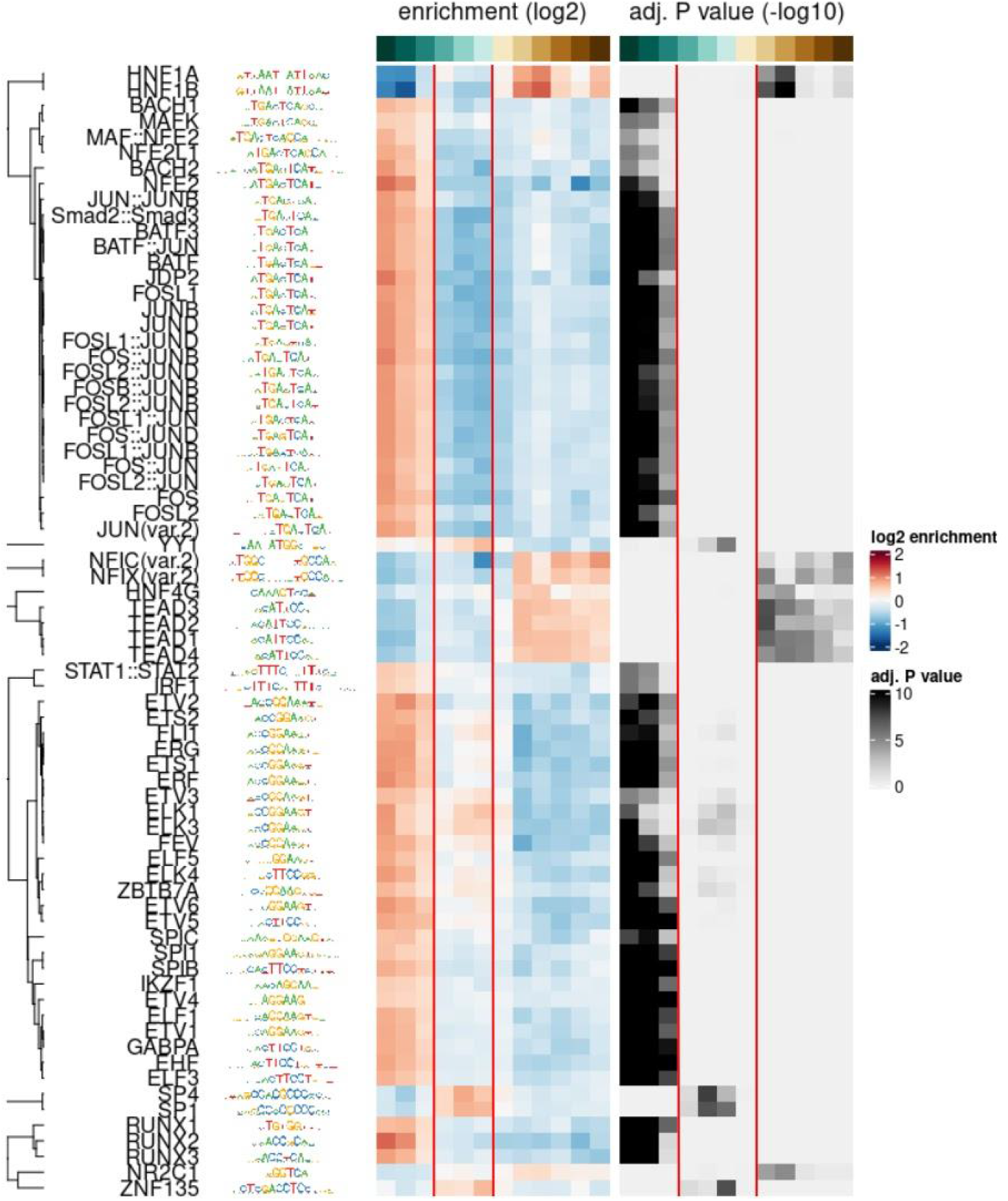
Methylation sites that correlate to IF are enriched for specific TF binding sites. TF enrichment and p-values of sequences around CpG sites that positively correlate with IF (left), do not correlate with IF (middle; between the red lines) and negatively correlate to IF (right).

### Kidney-specific methylation patterns are lost with kidney disease progression

Tissue-specific differentially methylated regions (tsDMRs) can have tissue-specific roles (Irizarry et al. 2009; Davies et al. 2012). After noticing the unique methylation pattern of cg10832035, we sought to test if this is a general phenomenon. Using the NGDC-CNCB (Xiong et al. 2021) dataset, we identified 427 methylation sites that exhibit a unique methylation pattern in the kidney (**Supp. Table 7**). Of these sites, 68% were in the top 10% (r>0.47; p<4**×** 10^−5^) of sites correlated with IF. Removing CpG sites with significant missing data increased this percentage to 72% (208 of 289; expected: 29; p=10^−322^; **Supp. Table 7**). Thus, the overwhelming majority of kidney unique methylation sites change their methylation pattern with functional decline.

### Function decline is accompanied by a drift towards the common methylation pattern

Cg10832035 is partially methylated in the kidney, but almost completely methylated in every other tissue examined (**Fig. 3E**). Indeed, we found this to be the common case. Of the 289 sites analyzed, 253 (88%) had a lower methylation level in the kidney as compared to the rest of the tissues, while only 36 (12%) had a higher methylation level. Upon kidney functional decline, cg10832035 becomes more methylated, trending towards the common methylation state of all other tissues examined. More generally, this raises three distinct possibilities, that upon functional decline (i) methylation levels increase (ii) methylation levels trend towards the common form (iii) methylation may increase or decrease independently of the common form. To distinguish between these three possibilities, we examined the levels of methylation in kidneys with different levels of fibrosis. For all 289 sites, methylation level trended toward the common form in the body (p=10^−87^), increasing in the 253 sites that are less methylated in the kidney compared to other tissues (**Fig. 5A**) and decreasing in the 36 sites that are more methylated in the kidney (**Fig. 5B**). Moreover, there was a positive correlation between the “uniqueness” of each site (as measured by the distance from all other tissues) and the correlation of the site with IF score (**Fig. 5C**; r=0.95 p<10^−148^). We concluded that kidney functional decline was associated with an erosion of the unique methylation pattern of the kidney.

**Figure 5:**
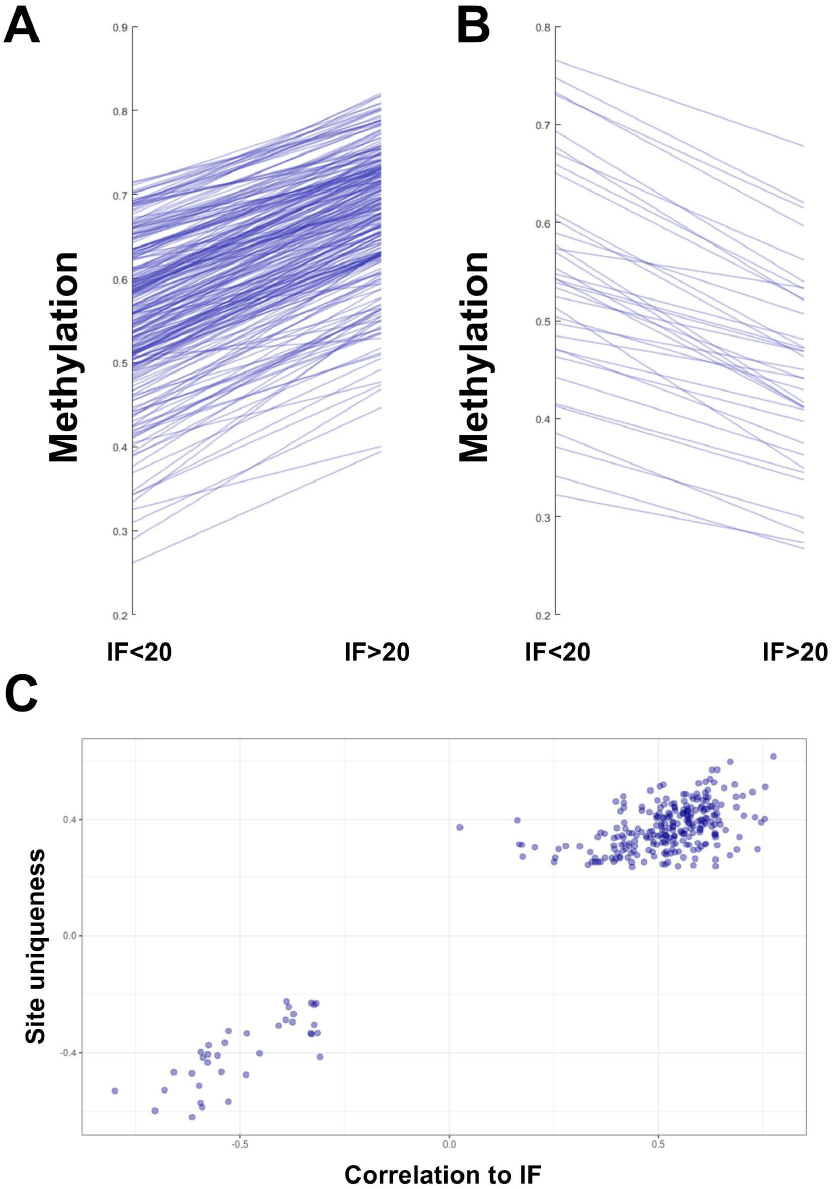
Methylation levels regress to the common baseline upon CKD. **A**. Methylation levels for the 253 sites undermethylated in the kidney, in individuals with IF<20 and >20. **B**. Methylation levels for the 36 sites overmethylated in the kidney, in individuals with IF<20 and >20. **C**. Site uniqueness, as measured by the distance from other tissues, and the corresponding correlation to IF.

### Characterization of kidney-specific methylation sites

Methylation patterns may be influenced by protein-DNA interaction, which in turn are guided by the primary DNA sequence. To identify motifs that may help establish or maintain these unique methylation patterns, we extracted the sequences surrounding the 253 and 36 identified sites, and looked for motif enrichment. *De novo* motif discovery, using random CpG sites as background, identified multiple sequences showing strong enrichment (**Supp. Fig. 2**). Not surprisingly, these included significant enrichment for motifs identified in CpGs that correlated with CKD progression. To identify sequences that potentially contribute to the unique methylation pattern, while removing general CKD background, we repeated this analysis using as background CpG sites that correlate to IF, but do not have a unique methylation pattern. Again, multiple motifs were identified, including HNF1 and HIF2a (**Supp Fig. 3**). However, the 253 sites identified significantly differ from both backgrounds. As most sites are almost fully methylated in all but one tissue, they may differ in GC content from the background. Indeed, the average GC percentage of both backgrounds, as well as the methylated-in-kidney sites, was ~55%. By contrast, the average GC content around the 253 sites (predominantly methylated in the body, with an exception in the kidney) was 45%. Thus, we compared these sites showing a unique methylation pattern, but in other tissues. Again, we saw a significant enrichment for specific TFs, including HNF1 (**Supp. Fig. 4**).

### Expression change of genes associated with kidney-specific methylation patterns

Changes in methylation levels are often associated with changes in gene expression. Depending on the genomic context, these changes can be causal or non-causal, and can involve an upregulation or downregulation of the associated gene (Dhar et al. 2021; Moore et al. 2013). We tested if changes to kidney-specific methylation patterns are associated with gene expression changes. To achieve this aim, we identified genes associated with the identified CpGs (**Supp. Table 7**), and tested for changes in mRNA levels in kidney biopsies of CKD patients and controls (Nakagawa et al. 2015). Of the 250 genes tested, 210 (84%) showed a statistically significant (p<0.05) difference between CKD and control (**Supp. Table 8**). Of these ~90% were upregulated in CKD (**Fig. 6**; **Supp. Table 8**). We concluded that methylation in these sites correlates with gene expression, likely by reflecting local chromatin structure.

**Figure 6:**
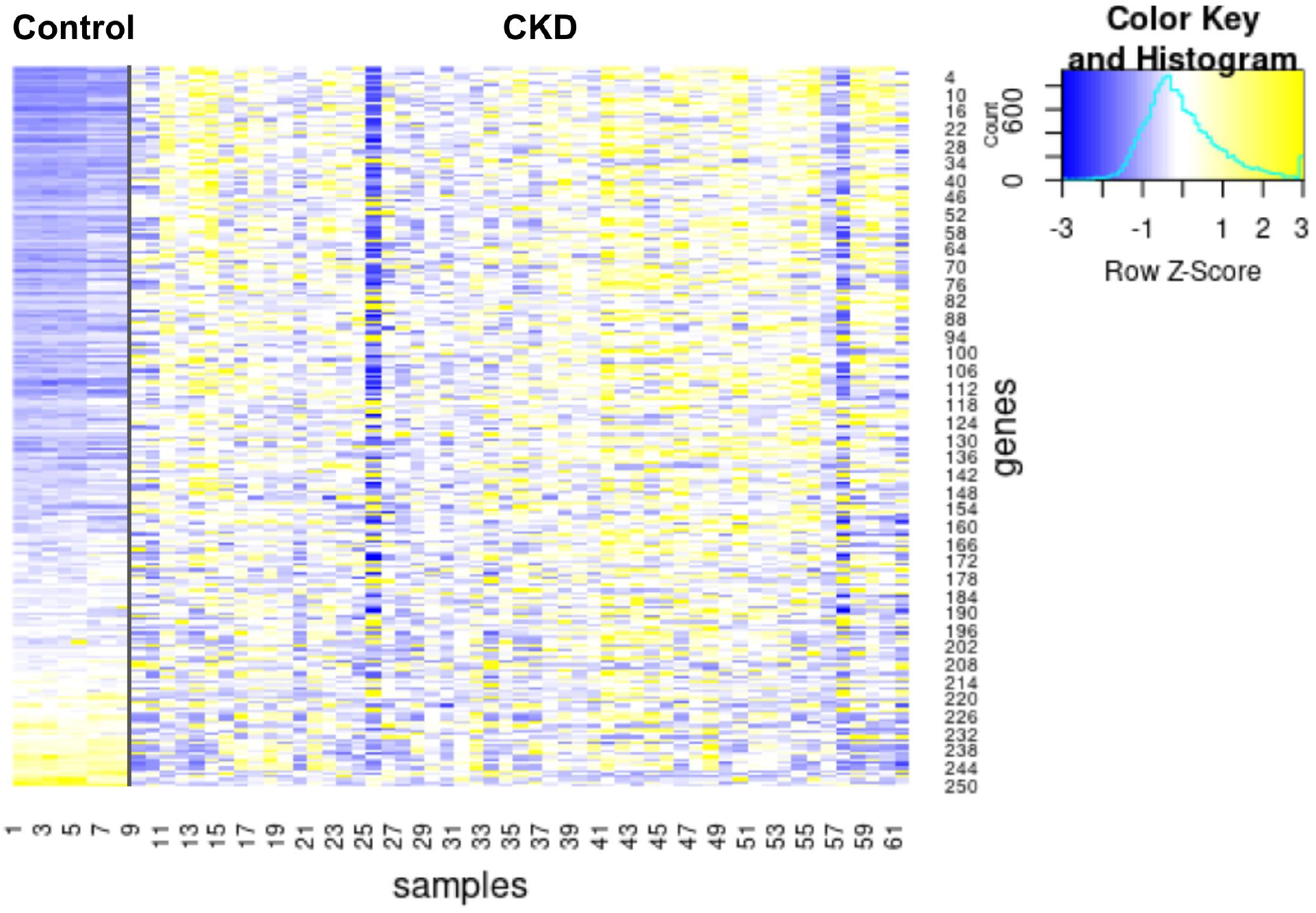
Genes associated with uniquely methylated sites change in expression in CKD. Expression levels of all (250) genes associated with uniquely methylated sites. Gene expression from the Nakagawa CKD Kidney dataset, with the discovery and validation cohorts united.

### Methylation sites negatively correlated with IF are linked to oncogenes and developmental genes

We clustered the 36 methylation sites negatively correlated with IF into 15 genomic regions which mapped to 13 genes (**Supp. Table 9**). Of these 13 genes, seven are known as renal cancer prognostic genes (overexpression), five take role in kidney differentiation and development, two of which overlap with cancer prognostic genes, and one functions as a tumor suppressor. Kidney fibrosis is known to correlate with CKD and subsequently with renal cancer (Lowrance et al. 2014; Christensson et al. 2013). SPAG5, MCF2L, CCDC64, PDLIM4, EMX1, P4HA2, VIM and GRAMD1B methylation declined for IF>20, which might indicate oncogenesis in the fibrotic tissue. Interestingly, CKD is a well-established risk factor for renal cancer (Lowrance et al. 2014; Christensson et al. 2013). Although kidney inflammation (Silverstein 2009) and relative immunodeficiency (Giacchino et al. 1982) have been suggested to play a role in CKD-related carcinogenesis, the exact molecular link has yet to be defined. Notably, although CKD results in a generalized inflammatory state (Shlipak et al. 2003), the risk of cancer in CKD has been shown to be limited to the kidney and not other organs (Silverstein 2009), implying that it likely arises due to kidney-specific changes. The described methylation changes, which are highly kidney-specific, may thus account for at least some of the enhanced cancer risk among CKD patients.

### Epigenetic information loss correlates with IF and eGFR

The discovery that epigenetic information is lost in a specific subset of methylation sites opens us the question: can a pathological state be predicted from methylation data alone, without any other prior knowledge. To answer this question, we implemented a naive information loss score. For each of the 289 unique methylation sites, discovered in the NGDC-CNCB dataset, the kidney median and standard deviation were calculated. Next, for each individual, the number of sites that deviate from the tissue median towards the levels in other tissues, by at least two standard deviations, was calculated (**Fig. 7**). This score naively represents the loss of epigenetic information in a specific kidney sample. The information loss score showed a r=0.82 correlation with IF (p<10^−18^) and r=−0.54 with eGFR (p<10^−6^). Thus, a pathological condition can be probabilistically observed merely by noting the loss of epigenetic information, without any prior knowledge of the disease to affected tissue.

**Figure 7:**
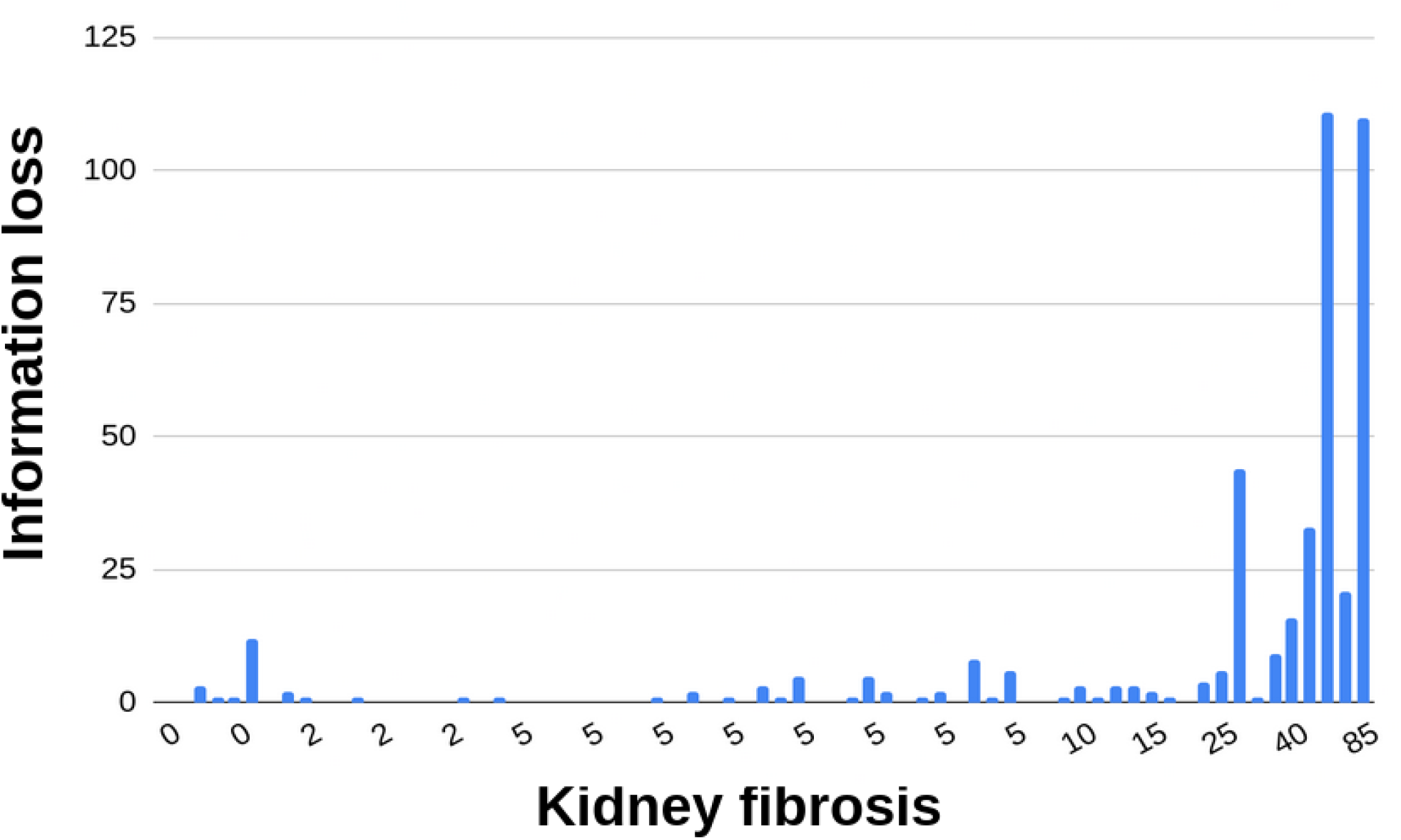
Epigenetic information loss correlates with IF. Epigenetic information loss score for each individual of the GSE50874 dataset (y-axis) along with the corresponding IF score (x-axis).

## Discussion

### Kidney state is captured by specific CpG methylations

Analysis of the kidney methylome revealed that multiple kidney pathologies are captured by distinct epigenetic clocks (**Fig. 1,2**; **Supp. Fig. 1**). DNAmB2M and DNAmPAI1 had a particularly good correlation with IF in males (r=0.75 and 0.58 respectively). DNAmB2M clock was built based on Beta-2 Microglobulin levels (Lu et al. 2019). Increased B2M levels in the blood, and by contrast low levels in the urine, are indicative of glomerular dysfunction (Argyropoulos et al. 2017; Sedighi et al. 2015; X et al. 2014), suggesting a possible explanation. Of note, DNAmB2M and DNAmPAI1 showed relatively low correlation amongst themselves (r=−0.30 in males and −0.14 in females), indicating that an improved estimator can be generated to facilitate more accurate estimation of kidney function across both sexes. Indeed, a global analysis identified multiple methylation sites, showing strong correlations for both sexes. A future increase in the number of samples will enable the construction of new epigenetic clocks, capturing kidney function decline with greater precision.

Next, we identified individual CpG sites that correlated with IF or eGFR. As expected, IF and eGFR associated CpGs showed a significant overlap. On average, IF resulted in stronger correlations than eGFR both for methylation clocks and specific CpGs. This is also expected, as eGFR is an inherently noisy measurement that does not capture the kidney state as accurately as IF (Levey et al. 2009). As expected, these results greatly overlapped with Bontha et al. 2017. By contrast, only a small overlap of 21 of 69 CpG sites was observed when compared to blood methylation markers of CKD (Meta-analyses identify DNA methylation associated with kidney function and damage). We concluded that the physiological state of the kidney is reflected by the methylation levels at multiple sites.

### Tissue-specific methylation patterns are lost with functional decline

Of all sites passing our rigorous threshold for tissue-specificity, 72% have a methylation pattern that correlates with IF (**Supp. Table 7**). This is likely an underestimation, as the arbitrary cutoff of 10% (p<4**×** 10^−5^) is guided by the relatively small dataset used to identify correlated CpGs. The loss of unique epigenetic signature with functional decline appears to be nearly universal in the kidney. Loss of epigenetic information has been suggested to accompany and potentially drive cellular aging (Lu et al. 2020; Imai & Kitano 1998). These findings support this hypothesis, and point that some of the loss may correlate better to functional decline or “biological age” than to chronological age. Moreover, tissue-specific methylation sites appear to be particularly vulnerable to such loss, as most uniquely methylated sites were associated with functional decline, compared to much smaller numbers among all sites. This exceptionally high occurrence leads us to speculate maintenance of these unique sites may serve as a hallmark of functional tissue. Future studies will test if this phenomenon also applies to other tissues, as well as functional decline that does not involve fibrosis.

### Uniquely methylated sites revert to the average

Interestingly, in every single uniquely methylated site tested, the functional decline was accompanied by shift in methylation levels towards the levels common in other tissues. The most plausible explanation for this would be that a default pattern exists in these sites, and a deviation from this default requires active maintenance. Primary DNA sequence is a key driver of local chromatin architecture. The unique methylation patterns could be established and maintained by specific DNA motifs. We tested for DNA motifs that could explain these patterns. While no single motif could explain this unique pattern, we have identified several candidates, enriched in these sites. *De novo* motif identification discovered not only motifs depleted near unrelated CpG sites, but also motifs present in uniquely methylated sites correlating to IF, but depleted near non-uniquely methylated sites that do correlate with IF.

While this work focused only on kidney-specific methylation patterns, we did observe multiple sites that were shared with one or a few other tissues and were excluded from our lists. It is likely that these findings can be greatly expanded, both by testing sites that show a distinct methylation pattern in other tissues, and by analyzing sites that show a distinct methylation pattern in a subset of tissues.

### Gene expression correlates with changes in methylation

DNA methylation can reflect aspects of local chromatin structure, thus only correlating to gene expression. Alternatively, DNA methylation alters the binding for some TF (Héberlé & Bardet 2019), affecting gene expression. We identified multiple genes, associated with CKD-correlated CpG sites, that change in expression in CKD (**Fig. 6**; **Supp. Table 8**). CKD-correlated CpG sites are enriched for JunD TF binding sites. DNA methylation inhibits JunD binding (Mann et al. 2013; Spruijt et al. 2013), thus providing a potential causal mechanism by which methylation at these sites regulates gene expression.

## Materials and methods

### Epigenetics clocks and CpG site correlations

Epigenetic estimators were calculated using the DNA Methylation Age Calculator (Horvath 2013) for the 85 Illumina HumanMethylation450 BeadChip array samples for which full pathological data was available (GEO GSE50874). Samples that generated the warnings “meanMethBySample <0.25” were removed and the data was re-analysed (**Supp. Table 1**), resulting in a very mild increase in most correlations. Next, samples separated into males and females and analyzed again (**Supp. Table 2-3**). All CpG sites were scored by Pearson correlation (r) to eGFR and IF. For each r, t=r√(n-2) / √(1-r^2^) was calculated, along with the corresponding p-value. Age and sex were were excluded as confounders by testing for association with eGFR and IF both the GSE50874 and the NGDC-CNCB dataset.

### Identification of uniquely methylated sites

The National Genomics Data Center, China National Center for Bioinformation (NGDC-CNCB; Xiong et al. 2021) dataset contains tissue-level methylation data across multiple partially annotated individuals. It was used to infer tissue-specific methylation levels. Uniquely methylated sites were defined as such that the average methylation level in kidney is lower by at least 0.2 than the 5% quantile level of all other samples in the NGDC-CNCB dataset, or higher by at least 0.2 than the 95% quantile level of all other samples. This distinguished methylation sites that out of the 28 tissues in the dataset, have a unique signature in the kidney, but did not exclude samples where a very small minority of samples leaks throughout the entire range. Next, CpG sites with less than 2000 values were removed. These sites were identified using custom R and Python scripts.

### Motif Enrichment

Motif enrichment analysis for CKD-correlated CpG sites was done using MonaLisa (Dania Machlab, Lukas Burger, Charlotte Soneson, Michael Stadler 2021), using 200bp of sequence from each side of each CpG site. As a control, CpG sites showing no correlation were inserted between the positively and negatively correlated sites. *De Novo* identification of motifs near uniquely methylated sites was done using Homer (Heinz et al. 2010) on 500bp of sequence from each side of each CpG site. These were compared to randomly selected CpG sites, or IF correlated randomly selected CpG sites.

### Gene expression analysis

Genes associated with all uniquely methylated sites CpG sites were extracted from the HumanMethylation450 v1.2 Manifest File (Illumina, USA). Gene expression levels in CKD and control kidney biopsies from the Nakagawa CKD Kidney dataset (Nakagawa et al. 2015) were extracted using NephroSeq.

## Author contribution

DZB and NS designed the experiments. NS performed the experiments with inputs form NM, PB, OP and DZB. NM performed the monaLisa analysis. DZB wrote the manuscript with inputs form NS, PB and OP.

## Funding

The work was supported by the Israeli Science Foundation (grants 654/20 and 632/20 to DZB).

## Conflict of interest

The authors declare no conflict of interest.

## Data availability

Data derived from public domain resources. New analysis data are available in the supplementary material.

## Acknowledgments

We thank the Bar lab members, Dr. Maayan Gal and Prof. Yosef Gruenbaum for critical reading of the manuscript, useful suggestions and comments.

## Supplementary figures

**Supplementary figure 1:**
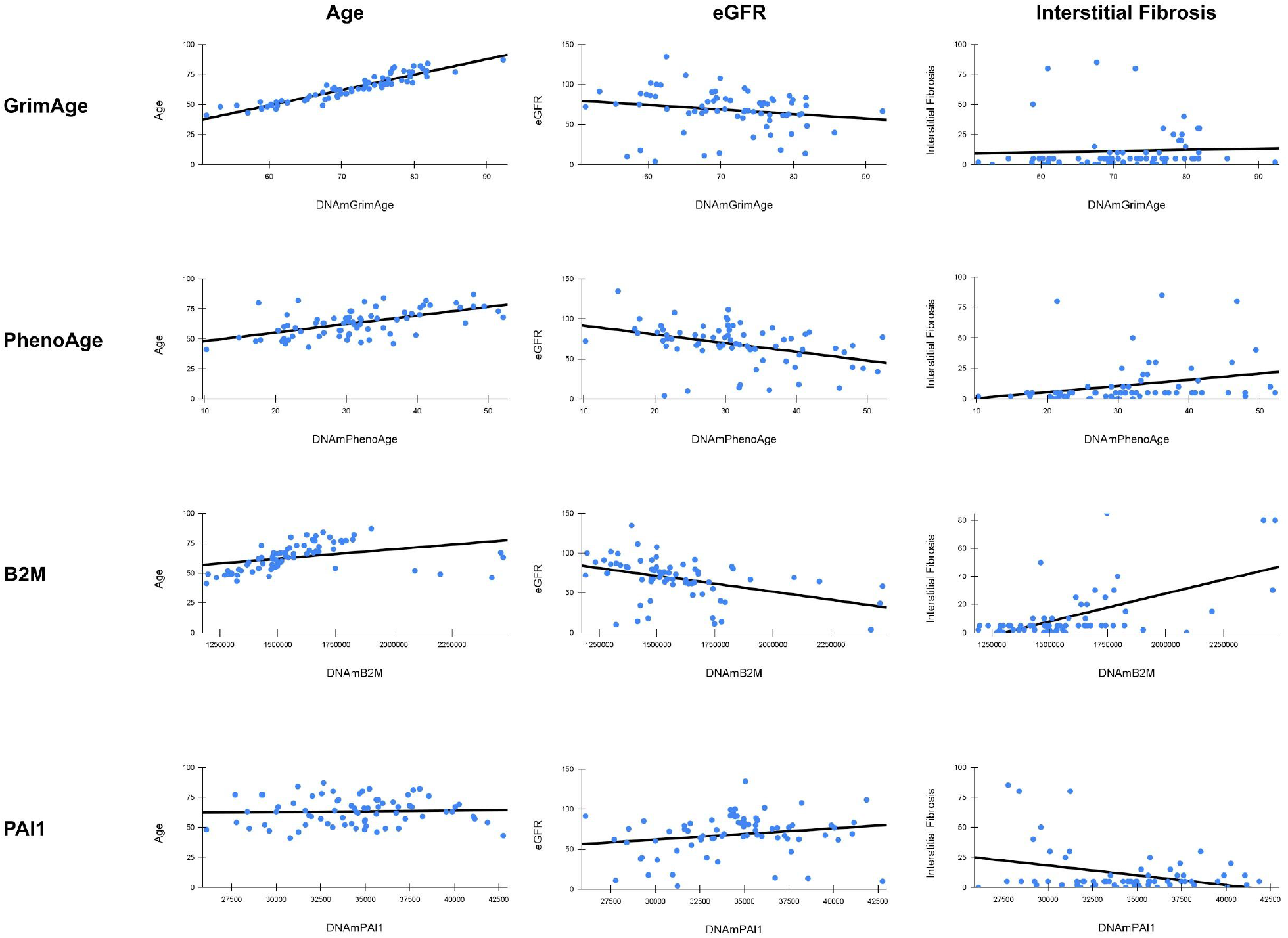
Methylation clocks correlate with kidney state. Correlation of age, eGFR and IF with four epigenetic clocks: GrimAge, PhenoAge, B2M and PAI1.

**Supplementary figure 2:**
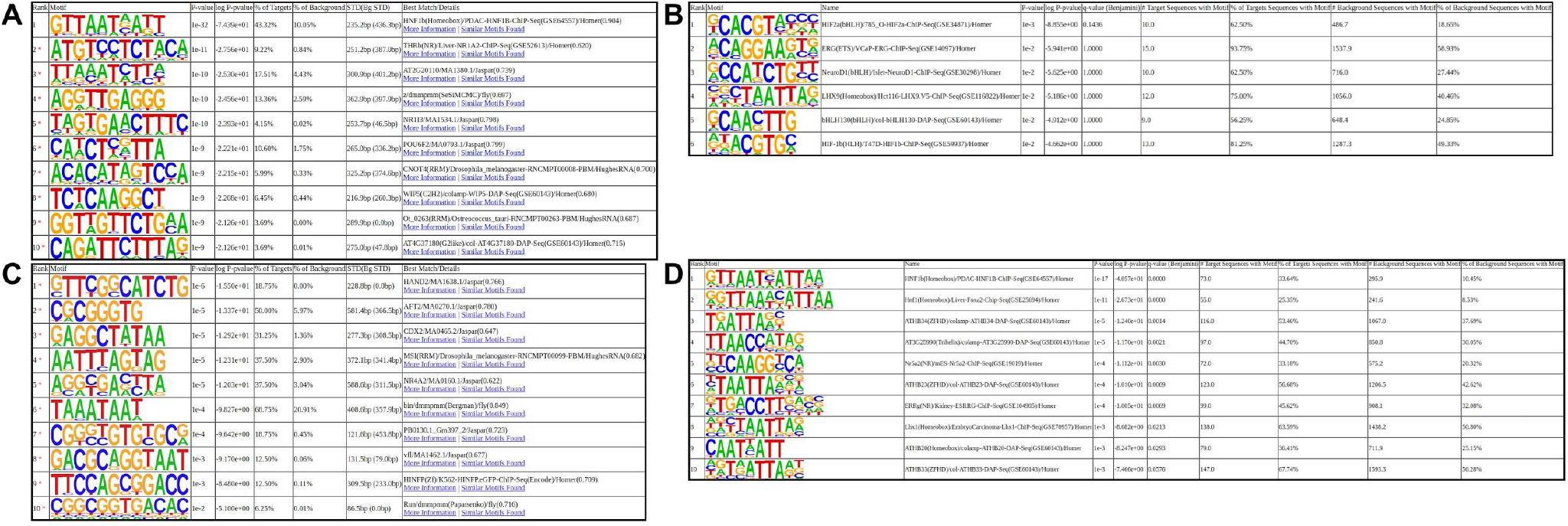
Motif enrichment in uniquely methylated sites vs. random sites. De novo (**A**) and known (**B**) motif enrichment in sites undermethylated in the kidney. De novo (**C**) and known (**D**) motif enrichment in sites overmethylated in the kidney. Up to top 10 results are displayed. * - possible false positive.

**Supplementary figure 3:**
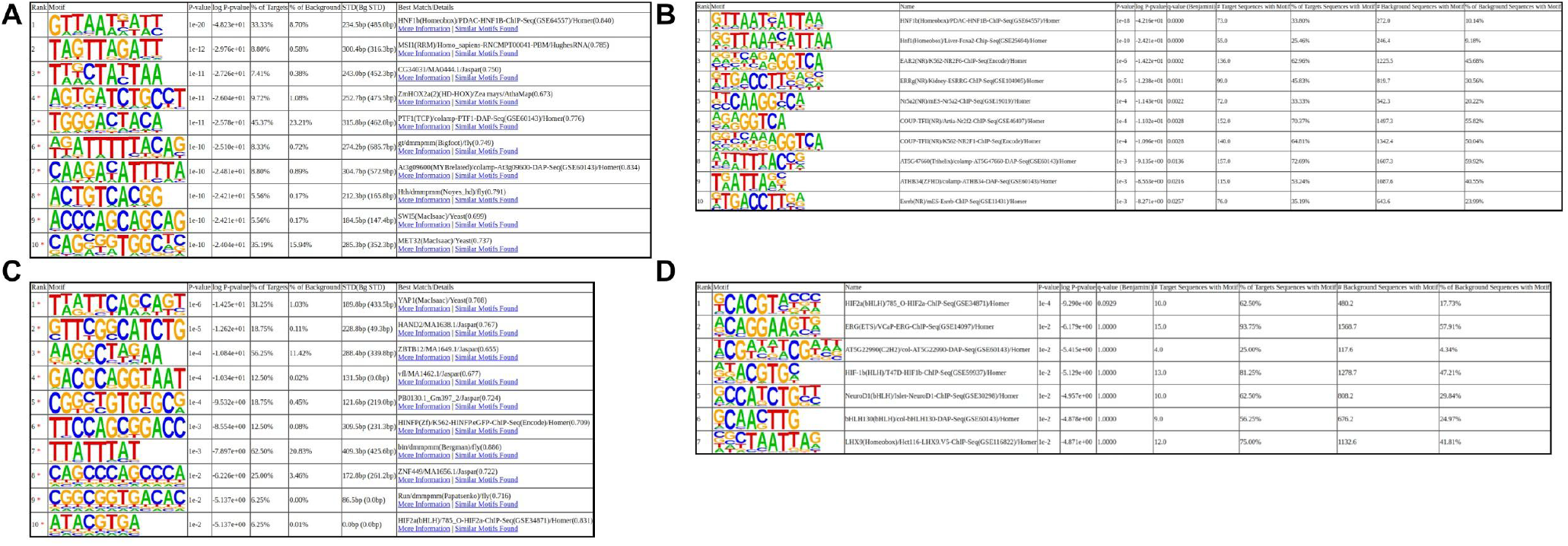
Motif enrichment in uniquely methylated sites vs. IF-correlated sites. De novo (**A**) and known (**B**) motif enrichment in sites undermethylated in the kidney. De novo (**C**) and known (**D**) motif enrichment in sites overmethylated in the kidney. Up to top 10 results are displayed. * - possible false positive.

**Supplementary figure 4:**
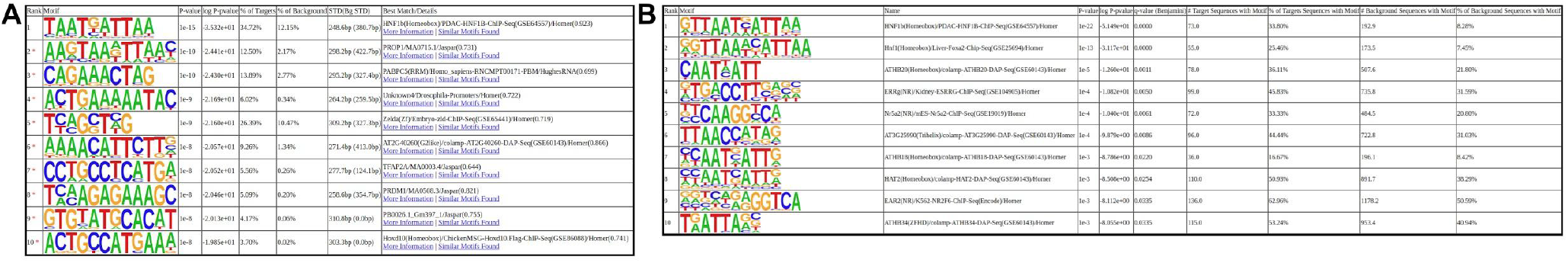
Motif enrichment in sites uniquely methylated in kidney vs. other tissues. De novo (**A**) and known (**B**) motif enrichment in sites undermethylated in the kidney. Up to top 10 results are displayed. * - possible false positive.

## Notes

### Competing Interest Statement

The authors have declared no competing interest.

### Summary of Updates

This revision includes text clarifications, analysis of putative confounders, as well as figure 7, showing an epigenetic information loss score and its correlation to kidney fibrosis.

